# Rate of Fixation of Rare Variants in a Population

**DOI:** 10.1101/123232

**Authors:** Bhavin S. Khatri

**Affiliations:** The Francis Crick Institute, 1 Midland Road, London, NW1 1AT, U.K.; Division of Infection & Immunity, University College London, London, WC1E 6BT, U.K.

## Abstract

The process of molecular evolution has been dominated by the Kimura paradigm for nearly 60 years; mutations arise at a certain rate in the population and they go to fixation with a probability given by Kimura’s classic formula, which assumes there are no further mutations that interfere with the fixation process. An alternative view is that rare variants exist in the population in a mutation-drift-selection balance and rise to fixation through a combination of chance (genetic drift), selection and mutation. When mutations increase in strength, but still in the weak regime, we would expect the Kimura rate approximation to be an overestimate, as a rare variant which grows in frequency will suffer a greater backward flux of mutations, slowing progress to fixation. However, to date calculating important quantities for a general model of selection and mutation, like the rate of fixation of these rare variants has not been tractable in the conventional diffusion approximation of population genetics. Here, we use Fisher’s angular transformation to convert the frequency-dependent diffusion inherent in population genetics to simple diffusion in an effective potential, which describes the forces of selection, drift and mutation. Once this potential is defined it is simple to show that the mean first passage time is given by a double integral which relate to populations at the barrier. Exact numerical integration shows excellent agreement with discrete Wright-Fisher simulations, which do show a slowing down of the fixation of mutants at higher mutation rates and for strong positive selection, compared to the Kimura prediction. We then seek a closed-form analytical expression for the rate of fixation of mutants, by adapting Kramer’s approximation for the mean first passage time. This overall gives an accurate approximation, but however, does not improve on the Kimura rate.

## I. INTRODUCTION

The probability of fixation of a mutant in a wild-type population is a key quantity in population genetics; given some initial frequency *x*_0_ in a population of finite size *N*, what is the probability that the frequency of this mutant variant reaches *x* = 1. In the simple case with no mutations, Kimura calculated his famous equation^1^, which describes a probability of fixation which has a saturating form, in the diffusion limit of the Wright-Fisher model. This has formed the basis of understanding the substitution process in studies of molecular evolution in the weak mutation regime, where mutations arise do novo and fix with a probability given by Kimura’s equation; the overall rate of substitution of different mutants is simply the per individual mutation rate multiplied by the population size and Kimura’s fixation probability.

However, in reality when mutations are of sufficient strength, but still in the weak regime (*N_μ_* < 1), we would expect even when one variant is nominally “fixed”, there will be polymorphisms that co-exist at low frequency in the population. Further, as a variant goes towards fixation, back mutations should retard its progress, causing a slowing of the rate of fixation, compared to the Kimura rate. If mutations are included in the diffusion approximation of a Wright Fisher process, it is not possible to calculate the mean time to fixation in simple closed-form. A key difficultly of the diffusion approximation is that the effective diffusion constant for variant frequencies, depends on the frequency of the variant. When a variant is close to fixation or loss, diffusion slows down compared to intermediate frequencies since the variance in the change in variant frequency has a characteristic binomial form. Fisher found a transformation to an angular frequency, where diffusion is independent of angular frequency, which, however, comes at the cost of introducing an effective non-linear potential that describes the flux of diffusers to the boundaries^2^. This non-linearity makes any exact calculations of the dynamics difficult, but Khatri^2^, developed a heuristic method to find accurate asymptotic Gaussian Greens functions in the short-time limit. Here we show that given this potential, a standard technique can be used to calculate the MFPT, where it is given as a double intergral over the barrier and well populations. When compared to discrete Wright-Fisher simulations, numerical evaluation of this integral gives very accurate estimates of the MFPT and confirms that variants at higher mutation rates have their fixation rate retarded, compared to the Kimura rate. We also calculate a Kramers-type approximation^3^, which is classically used for the MFPT of chemical reactions, where the potential energy barrier impeding a reaction is large; comparing to the discrete Wright-Fisher simulations we find the calculation is accurate, but does not improve on the Kimura rate. In addition, the Kramers approximation, as for Kimura theory, fails to predict the reduction of fixation rate at higher mutation rates and strong positive selection.

## II. FISHER’S ANGULAR TRANSFORMATION FOR 2-VARIANTS

In the diffusion approximation the stochastic dynamics of gene frequency *x* (= *n*/*N*, where *n* is the number of copies of the mutant variant *a*_1_ and *N* the total population) is given by

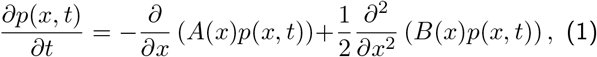

where *p*(*x, t*) is the probability density of gene frequency and *A*(*x*) is the mean change in gene frequency per generation and *B*(*x*) is the variance of gene frequency change per generation - we assume these only have a time-dependence though *x*(*t*). This is the forward Fokker-Planck equation whose solutions represent a progression forward though time given an initial condition *p*(*x*, 0) = *δ*(*x* − *x*_0_) - i.e. we know the initial gene frequency at time zero. For selection and arbitrary mutation:

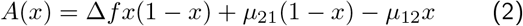

where Δ*f* = *f*(*a*_2_) − *f*(*a*_1_), so Δ*F* > 0 means selection favours variant *a*_2_, *μ*_12_ is the mutation rate from variant 1 → 2 and *μ*_21_ the mutation rate from variant 2 → 1 and

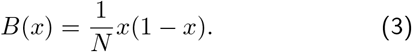

They key difficulty with Eqn.1 is that the variance of gene frequency change per generation depends on the frequency *x*. Fisher proposed a transformation to a different co-ordinate *θ*:

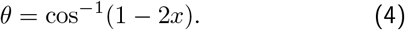

It is simple to show that the resultant Fokker-Planck equation is a diffusion equation in an effective potential

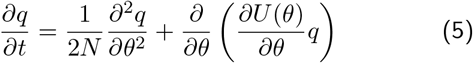

where the derivative of the effective potential in the above equation is given by,

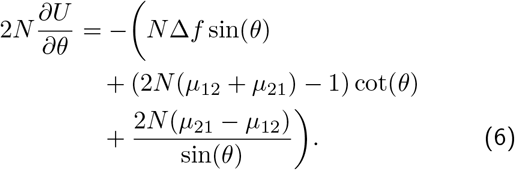

so that the potential is found on integration to be

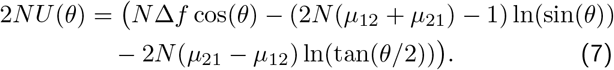

From Eqn.5 it is then simple to show that the equilibrium pdf is given by

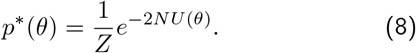

In Fig.1, we have a plot of the potential for 2*Nμ*_12_ = 0.1 and *μ*_21_ = 2*μ*_12_ for various values of 2*N*Δ*F*.

**FIG. 1.**
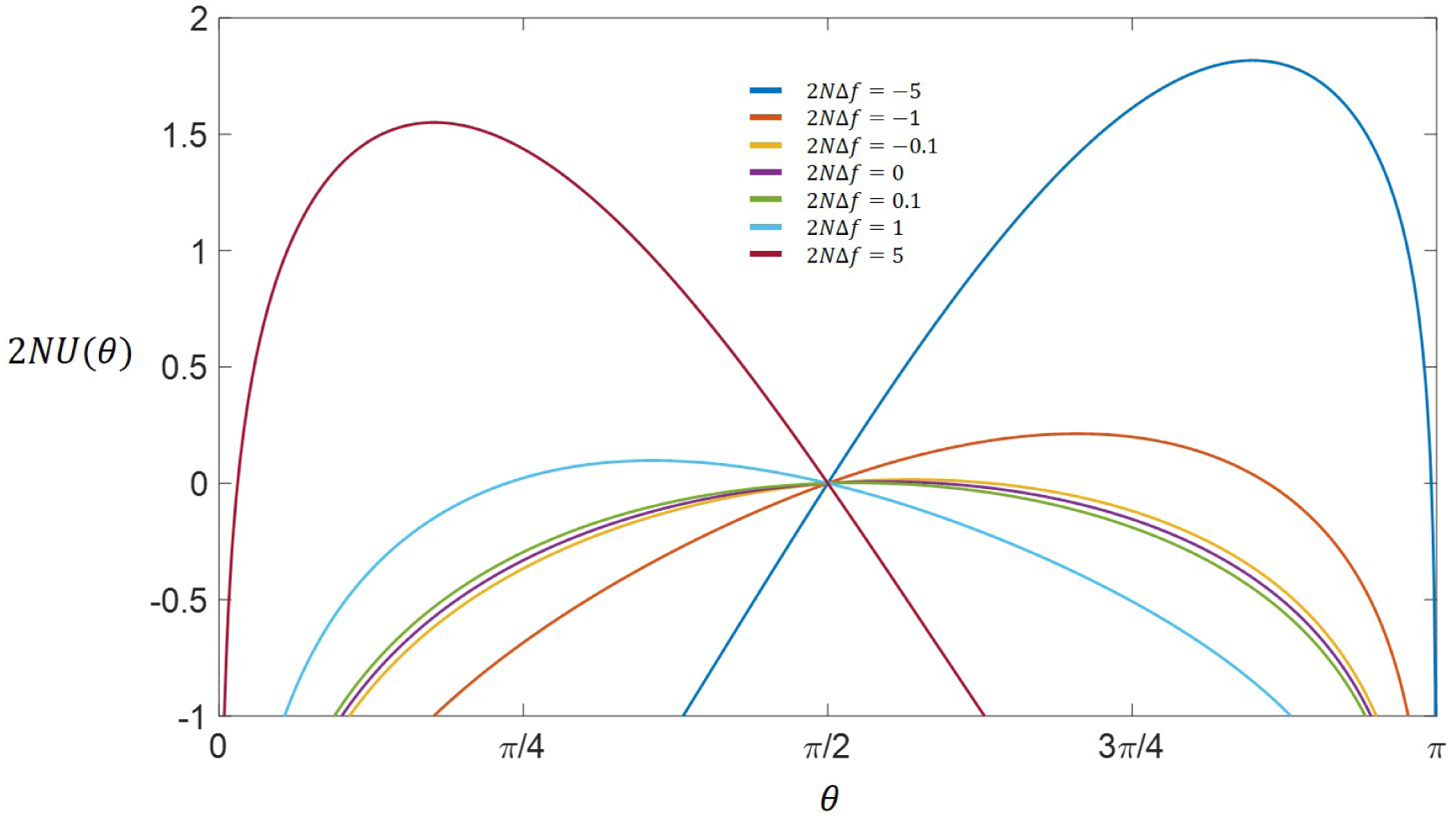
The effective potential *U*(*θ*) that arises after Fisher’s angular transformation Eqn.4, for 2*Nμ*_12_ = 0.1 and *μ*_21_ = 2*μ*_12_ for various values of 2*N*Δ*F*.

## III. MEAN FIRST PASSAGE TIME FOR 2-VARIANTS

The first passage time is the time it takes for a diffuser to *first* reach a boundary – this will have a distribution and we want to calculate it’s mean, which is the mean first passage time (MFPT). There are a number of ways to do this, one of which is to solve the corresponding backward Fokker-Planck equation^4,5^ for the mean first passage time for an initial frequency; however, under selection and mutation, no-known closed form solution is known. Here, we will assume that the initial frequency of the rare variant is not known, but that it is in quasi-equilibrium due to a mutation-selection-drift balance near *x* = 0 or *θ* = 0. After some period of time the rare variant will drift in frequency to a critical value, after which a combination of selection and mutation take the variant to fixation. The critical frequency *θ*\* will be given by the maximum in the effective potential *U*(*θ*). A method to solve such problems was developed by Kramers^3^. Here, the simplest exposition is to consider that we inject rare variants into the system at *θ* = 0, at a rate *J*, and then remove them when they reach fixation. From Eqn.5 the net flux in the system must equal to a constant, the number we inject per generation *J*:

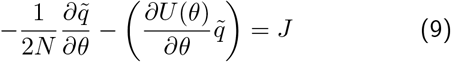

where 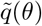 is the non-equilibrium *steady-state* distribution when there is a net flux through the system. Its integral is the number of diffusers in the system and so the MFPT *τ* simply obeys

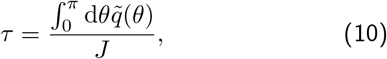

An expression for the steady-state distribution can be obtained by integrating Eqn.9:

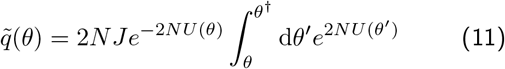

where *θ*^†^ is the angular frequency, which corresponds to fixation - as we shall see our answer will not be very critical on the exact value of *θ*^†^. Using Eqn.10 and 11, and swapping the order of integration, we find the MFPT is given by the following double integral:

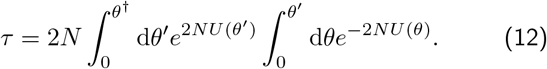

This expression is exact and can be numerically integrated for given values of Δ*f, N, μ*_12_, and *μ*_21_. However, the potential is singular at *θ* = 0 and *θ* = *π*, which requires a careful numerical integration scheme that essentially integrates a modified integrand, where the singularity is removed by choice of a function that has the same limiting form at the singular points and is itself is integrable over the region of interest. Using the limiting form lim_*θ*→0_{*e*^−2*NU*^} → *e*^−*N*Δ*f*^2^−2*N*Δ*μ*^θ^4*N*^*μ*_21_ – 1 as a function that is simple to integrate, we perform this numerical integration using a standard numerical routine in Matlab. The results are shown in Fig.2 and show excellent agreement with the MFPTs calculated from simulations of the discrete Wright-Fisher process.

**FIG. 2.**
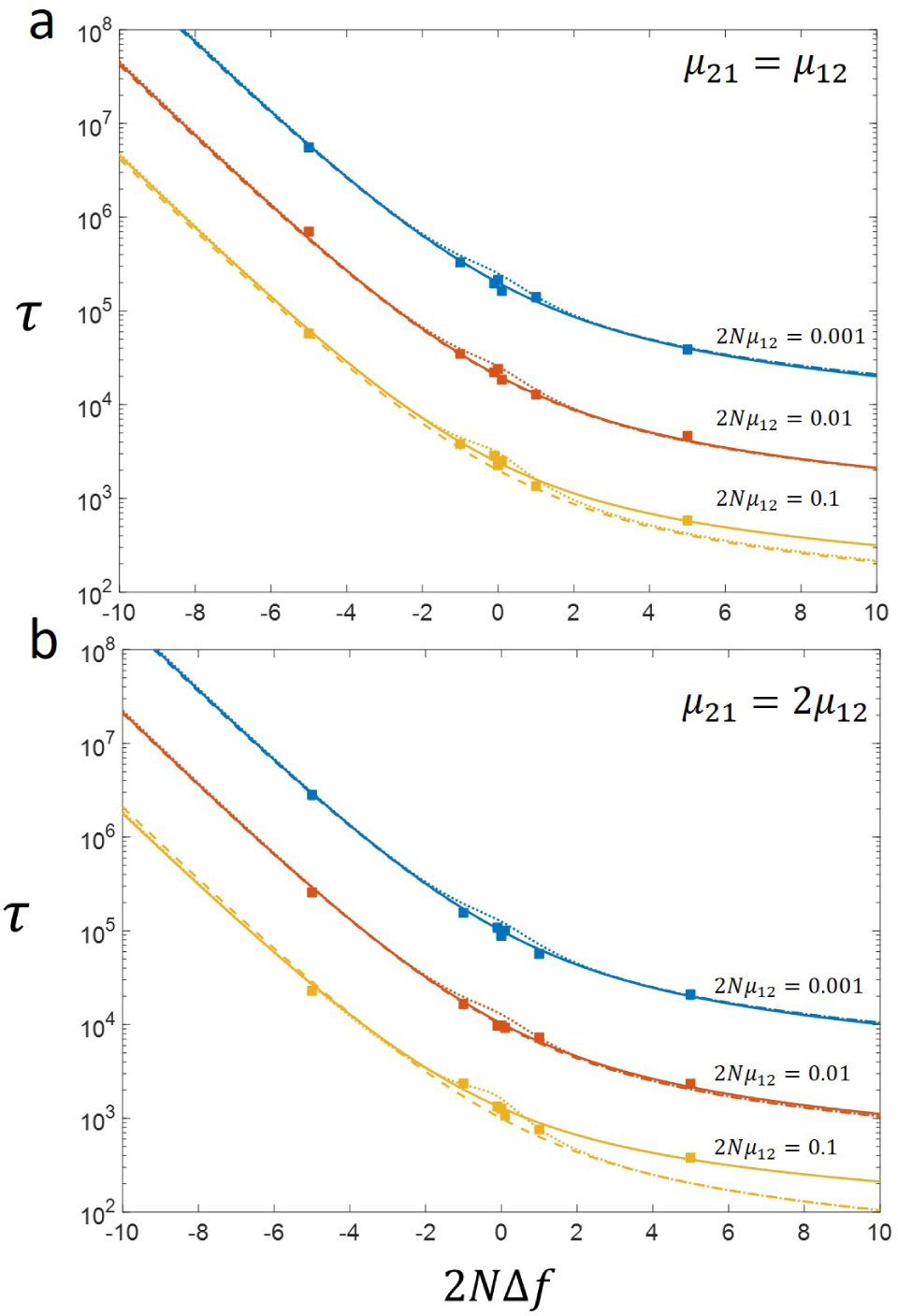
Comparison of discrete Wright Fisher simulations of MFPT (squares) to numerical integration of Eqn.14 (solid line) and Kimura’s approximation (dashed line) and Kramer’s method (dotted line). Top graph is for *μ*_12_ = *μ*_21_ and bottom graph is for *μ*_21_ = 2*μ*_12_.

We can also develop an approximation of the integral as follows, which is a modification of Kramer’s approximation^3^: the outer integral is of the form 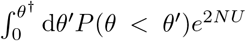 and so will be dominated by values of theta where *U* is maximum, which when mutation is weak, will correspond to the top of the barrier of *U* – at the barrier the function *P* (*θ* < *θ*′) (which comes from the inner integral above) will vary slowly as most of the density will be to the “left” of the barrier, so we can take this factor out of the integral, evaluated at the barrier *θ*\*, to give *τ* ≈ 2*NP*(*θ* < *θ*\*) ∫ d*θe*^2*NU*^. To evaluate *P*(*θ* < *θ*\*), we make the approximation that *θ* ≪ 1, 2*NU*(*θ*) ≈ *Ns*(1 − *θ*^2^/2) − (2*N*_*μ*_Σ__ − 1) ln(*θ*) − 2*N* Δ_*μ*_ ln(*θ*/2) (where *μ*_Σ_ = *μ*_12_ + *μ*_21_ and Δ*μ* = *μ*_21_ − *μ*_12_). The resultant integral that defines *P*(*θ* < *θ*\*) can then be rearranged by change of variable to give the definition of the lower incomplete gamma function 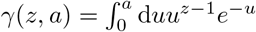, so that the inner integral is given by

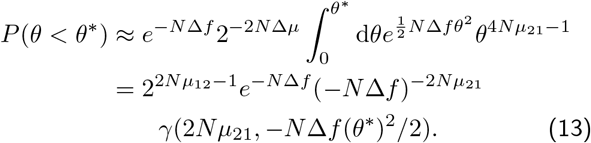

Next to evaluate the outer integral, we approximate the potential around the maximum of the barrier by a quadratic, so 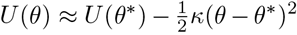, where *κ* = |d^2^*U*/d*θ*^2^|_*θ*=*θ*\*_ so that,

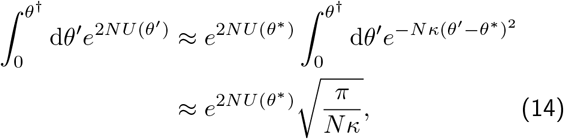

where we have extended the upper and lower limits to ±∞, so we can use the standard integral 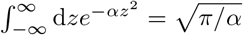.

This last approximation assumes that 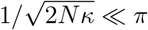, which as we can see from Fig.1 will not be very good for 2*N*Δ*f* ≪ 1; we will see in Fig.2 that if we replace this approximation for the barrier integral with a numerical integration, this regime is where this calculation is less accurate. The final expression for the MFPT is then

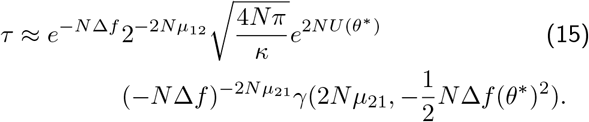

The rate of fixation of the rare variants is then *k* = 1/*τ*. The only task left is to calculate the position of the barrier, *θ*\* and the curvature at the barrier, *κ* = |d^2^*U*/d*θ*^2^|_*θ*=*θ*\*_. The first is done by solving for d*U*/d*θ* = 0, which gives a quadratic equation for cos *θ*\*, which has solution:

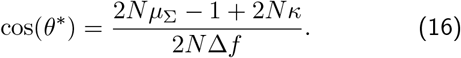

where *κ* is calculated directly from the second derivative of the potential as:

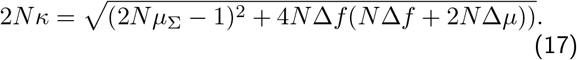

We can see that comparing to simulations in Fig.2, Eqn.15 is generally quite accurate, where this accuracy diminishes somewhat for 2*Nμ* = 0.1 and 2*N*Δ*f* = 5, where the calculation underestimates the MFPT. Finally, we compare this calculation to the Kimura rate:

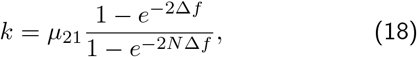

where *τ* = 1/*k* and we assume the net mutational input only depends on *μ*_21_. We see that the Kimura rate (dashed lines in Fig.2 very accurately predicts the mean first passage time, including the regime 2*N*Δ*f* ≪ 1, where Eqn.15, in contrast, is less accurate. However, for large and positive 2*N*Δ*f* the Kimura formula underestimates the MFPT, as also found with Eqn.15.

These results support the original hypothesis that when mutations begin to increase in strength (but still in the weak mutation regime *N_μ_* ≪ 1), we should expect to see that the Kimura approximation will overestimate the rate of fixation due to the retardation effect of back mutations as a variant approaches fixation; this is consistent with the observation that the effect is stronger when the difference in mutation rates Δ_*μ*_ > 0, such that back mutations are stronger. However, we find that this retardation effect is only significant when 2*N*Δ*f* > 1.

## IV. CONCLUSIONS

Calculating the rate of fixation of rare variants or polymorphisms in population is an important quantity to understand the rate of molecular evolution. Although, the Kimura approximation for the rate of fixation of mutants is widely popular, it ignores the effect of mutations, as a mutant rises to fixation. We should expect it to overestimate the rate of fixation for large mutation rates, due to an increasing flux of mutations back to the wildtype, as the mutant allele approaches fixation. To investigate this effect, we show that techniques from chemical reaction kinetics for calculating the rate of crossing a potential energy barrier, can be adapted to this canonical population genetics problem; first the frequency-dependent diffusion of the Wright-Fisher process is removed by Fisher’s angular transformation^6^, which results in simple diffusion or Brownian motion in an effective potential, which is analogous to the potential energy in a chemical reaction. The mean first passage time is then expressed as a double integral over the potential surface; we show that careful numerical integration of this integral gives excellent agreement with the MFPT calculated from discrete Wright-Fisher simulations. In comparison, we see the Kimura approximation does underestimate the MFPT, but only for larger mutation rates and for strong positive selection. The failure of the Kimura theory is likely due to the effect of back mutations retarding increase in frequency of the allele, as it approaches fixation; in support of this we find that the magnitude of the discrepancy between Kimura and Wright-Fisher simulations, increases as the ratio of the backward to forward mutation rate increases. We also evaluated this integral approximately using a modification of Kramers’ theory^3^, although overall this does not improve on Kimura’s theory.

In general, these results point to a new calculational technique, where Fisher’s transformation converts classic problems in population genetics to one of simple Brownian motion in a potential, which from the literature in physics and chemistry^4,7^ exist many approximate methods to find solutions.

